# An adversarial approach to guide the selection of preprocessing pipelines for ERP studies

**DOI:** 10.64898/2026.03.26.714586

**Authors:** Daniele Scanzi, Dylan A. Taylor, Katie A. McNair, Rohan O. C. King, Carley Braddock, Paul M. Corballis

**Affiliations:** School of Psychology, University of Auckland

**Keywords:** eeg, preprocessing, pipeline, ica

## Abstract

Electroencephalography (EEG) data are inherently contaminated by non-neuronal noise, including eye movements, muscle activity, cardiac signals, electrical interference, and technical issues such as poorly connected electrodes. Preprocessing to remove these artefacts is essential, yet the optimal method remains unclear due to the vast number of available techniques, their combinatorial use in pipelines, and adjustable parameters. Consequently, most studies adopt ad hoc preprocessing strategies based on dataset characteristics, study goals, and researcher expertise, with little justification for their choices. Such variability can influence downstream results, potentially determining whether effects are detected, and introduces risks of questionable analytical practices. Here, we present a method to objectively evaluate and compare preprocessing pipelines. Our approach uses realistically simulated signals injected into real EEG data as “ground truth”, enabling the assessment of a pipeline’s ability to remove noise without distorting neuronal signals. This evaluation is independent of the study’s main analyses, ensuring that pipeline selection does not bias results. By applying this procedure, researchers can select preprocessing strategies that maximize signal-to-noise ratio while maintaining the integrity of the neural signal, improving both reproducibility and interpretability of EEG studies. Although the data presented here focuses on processing and analysis most relevant for ERP research, the method can be flexibly expanded to other types of analyses or signals.

## 1 Introduction

Electroencephalography (EEG) data contain a mixture of neuronal activity, predominantly post-synaptic potentials of pyramidal neurons (Nunez & Srinivasan, 2006), and non-brain signals generally considered to be unrelated noise. Noise artefacts like heartbeat, blinks, muscle movements and electrical interference can conceal the signal generated by neurons and make EEG data analyses challenging. To address the issues introduced by noise, researchers usually preprocess their data before analysing them. Broadly defined, preprocessing comprises a suite of methods and algorithms designed to reduce or eliminate noise while preserving the neuronal signals.

Researchers have long sought to obtain clean data free from noise artefacts, employing techniques such as downsampling, filtering, averaging, automatic detection of noisy channels, and artefact removal via Principal Component Analysis (PCA), Independent Component Analysis (ICA), or Artefact Subspace Reconstruction (ASR) (Kothe & Makeig, 2013). At face value, having access to a rich preprocessing toolbox is excellent, as more tools could provide more denoising power. However, “with great power there must also come great responsibility”. The high number of available techniques increases the researcher’s degrees of freedom, making it challenging to compare results across studies. Moreover, no single method fully eliminates noise while preserving neural signals, complicating the evaluation of how preprocessing impacts downstream results (Luck & Kappenman, 2012).

Discussions within the EEG literature have revolved around determining optimal approaches for cleaning electrophysiological data. Some studies delve into specific aspects like comparing filter types and parameters (Acunzo et al., 2012; Rousselet, 2012; Tanner et al., 2015; Vanrullen, 2011; for a review and guidelines Widmann et al., 2015), evaluating referencing methods (Dong et al., 2019), or assessing the performance of various blind source separation algorithms (Barban et al., 2021; Delorme et al., 2012). In contrast, some researchers opt for a more comprehensive approach, focusing on contrasting entire pipelines rather than specific steps (Bailey, Biabani, et al., 2023; Bailey, Hill, et al., 2023; Delorme, 2023; Fló et al., 2022; Gabard-Durnam et al., 2018; Robbins et al., 2020); although note that these comparisons are usually conducted when presenting a new pipeline. Despite the wealth of investigation, a consensus on the optimal methods or pipelines remains elusive. Divergent conclusions across studies may be attributed to variations in analytic methodologies, dataset-specific nuances, or the testing of a different set of methods.

Regardless of each study’s specific focus, the literature on EEG data preprocessing commonly uses two approaches to evaluate the impacts of data-cleaning algorithms. The first approach employs simulated signals to quantify the extent of noise reduction after preprocessing (Barban et al., 2021; Grouiller et al., 2007; Hoffmann & Falkenstein, 2008; Makeig et al., 2000). The second approach uses actual EEG recordings, comparing results obtained by subjecting the data to different preprocessing steps (Robbins et al., 2020; Rousselet, 2012; Tanner et al., 2015). Both these approaches present limitations that warrant consideration when interpreting results. Using simulated noise artefacts could introduce a potential issue of “circularity”. If noise templates are generated using procedures or models similar to those that will be tested for their cleaning efficacy (Delorme et al., 2007; Dong et al., 2019), the results might be biased; a factor that needs to be considered during the analyses (Hoffmann & Falkenstein, 2008; Shackman et al., 2009). Additionally, accurately modelling EEG data, considering the time dynamics of both noise and neuronal signal, is a non-trivial task (Urigüen & Garcia-Zapirain, 2015). Using actual EEG recordings avoids the circularity problem but introduces the fundamental challenge of establishing a ground truth neuronal signal. We are always uncertain regarding which cleaning approach returns data that most closely resembles the true neuronal activity.

An often-overlooked aspect is that studies can only compare a subset of methods, limiting the generalizability of the results outside the sample of pipelines (or techniques) explicitly assessed. From a researcher’s viewpoint, this is problematic for two reasons. Firstly, there is no straightforward way to choose between two methods that have not been explicitly contrasted before. Secondly, most published pipelines allow users to modify default parameters or steps to suit their specific needs, further adding experimenters’ degrees of freedom. Thus, it is unclear whether the results presented in publications still hold even when different sets of parameters are used.

In this study, we propose a framework aimed at overcoming the limitations outlined earlier, offering researchers the ability to compare any pipeline of interest. Our method employs actual EEG recordings to evaluate the behaviour of different preprocessing methods against real noise. Crucially, it also introduces a “ground truth” signal to assess whether denoising algorithms preserve the integrity of the signal of interest. While preprocessing aims to eliminate noise, it would be counterproductive if the neuronal signal were inadvertently removed or altered. Although conceptually similar approaches have been presented in the past (semi-synthetic data: Barban et al. (2021); injected data: Chen et al. (2017); Delorme et al. (2007)), their primary intent was to investigate the effects of certain procedures and quantify the reduction of noise. Here, instead, we adopt a more general approach whose primary goal is to allow researchers to compare the efficacy of preprocessing pipelines when they need one to clean their data. We designed this method keeping four main objectives in mind:

1. *Flexibility*. Researchers should be able to compare any pipeline or variations of pipelines they are interested in. Providing this flexibility would help in making informed decisions even in those scenarios where established procedures need to be modified. We propose a framework that leaves to the individual researchers the choice and freedom to compare the methods that they prefer.
2. *Blindness to the real signal*. Researchers should test cleaning procedures using the same or similar (e.g. pilot data) datasets that they will analyse if they want to capture the idiosyncrasies of their data (system used, population tested, etc…). However, they should not select a cleaning procedure based on whether it produces statistically significant results in their study. We propose a method that can be applied to the data that will be analysed while remaining completely blind to the real signal of interest. Blindness is achieved by using an injected ground truth signal and simulating trials.
3. *Interpretability*. The metrics used to compare pipelines should be easily interpretable and understandable. Here, all our results are expressed as “the probability of a pipeline returning a better or equally good signal compared to another one”. We believe that an easy-to-interpret metric can be utilised by researchers at any career stage, from novices to seasoned professionals. Easily interpretable results allow individuals to capture the effects of a process, facilitating informed and reasoned decision-making. Indeed, in the development of this approach, we prioritised the interpretability of results over the applicability of a preprocessing method (that is, how easy a pipeline is to implement and use). This choice stems from the recognition that a pre-packaged preprocessing script, while running without errors, may yield problematic results. Without a sufficient grasp of the algorithms and their limitations, potential errors might go unnoticed. Consequently, we leave the selection and scripting of the pipelines to the researchers.
4. *Non-binarity of the results*. Debates in this field frequently revolve around identifying the best overall pipeline (Delorme, 2023). However, each study has its own peculiarities, and researchers often need to create their pipelines or alter, even slightly, existing ones. Consequently, it is unlikely that any single method can perform equally well across all possible scenarios (Urigüen & Garcia-Zapirain, 2015). We acknowledge this by proposing a framework that refrains from delivering a black-and-white statement on the best pipeline someone should use. By avoiding strong definitive statements, we emphasise the need for researchers to engage in informed decision-making, considering the unique aspects of their data, experimental set-up and expertise.

The code used to replicate the results presented here, as well as to run similar analyses on other datasets, is publicly available on the Open Science Framework (OSF) project directory https://osf.io/rg5ah/files/osfstorage. Original files are available upon request, as their size exceeds online storage limits. The MATLAB toolbox created to inject a simulated ground-truth signal in the data is freely available on Github https://github.com/d-scanzi/EEGinject. All procedures used in this study were approved by the University of Auckland Human Participants Ethics Committee (reference number UAHPEC26812).

### 1.1 Study 1

In the next sections, we describe the details of the procedure used to generate the dataset containing the ground truth signal and the analytical framework adopted to compare pipelines.

#### 1.1.1 Dataset Generation

##### EEG dataset

We recorded one of the authors (DS) completing an adapted version of the Simon task (Simon, 1969) with coloured stimuli (Craft & Simon, 1970; Hietanen & Pia, 1995). The details of the task are irrelevant to the current study and are reported in the supporting documentation online https://osf.io/rg5ah/files/osfstorage. Readers interested in the Simon task can refer to Cespón et al. (2020).

We opted for task-based EEG instead of resting state to ensure the dataset incorporated realistic noise elements such as blinks, eye movements, or muscle artefacts. Additionally, we wanted to use a dataset that reflected the characteristics of our data collection settings. Because the nature of the task, the EEG system used, and the recording procedures can all affect the type of noise introduced in the data, it is important to stress that the results presented might change if a different dataset is used. Researchers should test pipelines on their own data.

We collected approximately 35 minutes of data using a BrainVision Recorder with a gel-based active 64-channel Ag/AgCl “actiCap snap” system (Brain Products, Germany). Electrode placement adhered to the international 10-20 system covering the whole scalp. FCz acted as the online reference, and Fpz as the ground. Data was digitised at a sampling rate of 1000 Hz, and the impedance of each electrode was kept below 10 kΩ.

##### Ground truth signal generation

The ground truth signal was obtained through the following steps:

1. Generate a forward model
2. Generate a source signal
3. Project the signal to sensor space

We will address each step separately to provide a complete view of the process. Every step was conducted in MATLAB version 9.13.0.2049777 (R2022b) (The MathWorks Inc., Natick, MA).

##### Generate a forward model

A forward model describes how the current fields originated at the source level (brain) propagate through head tissues and are represented in the sensor space (electrodes) (Mosher et al., 1999). To generate the forward model, we used Brainstorm version 16-Nov-2022 (Tadel et al., 2011), which is documented and freely available for download online under the GNU General Public License (http://neuroimage.usc.edu/brainstorm).

We started by using a model of the inner skull, outer skull and scalp surfaces of the ICBM152 anatomy template (Fonov et al., 2009) provided as default in Brainstorm. We employed an anatomy template as we did not have access to a structural scan of the participant for which the EEG data was recorded (but see Guidance section for alternatives). The surfaces, represented as 3D triangular meshes, were extracted using the Brainstorm method with default parameters (vertices: scalp = 1922, outer skull = 1922, inner skull = 1922; skull thickness = 4mm). Electrodes were added by modifying the default 64-channel actiCap file co-registered to the ICBM152 template. Notably, we adjusted the template to match the electrodes in the recording cap, removing electrode FCz and Fpz (online reference and ground), and adding electrode AFz. The Boundary Element Method, implemented by OpenMEEG (Gramfort et al., 2010; Kybic et al., 2005), was then applied to obtain the forward model. We used the default relative isotropic conductivities (scalp = 1, outer skull = 0.0125, inner skull = 1), and set the source space to volume.

The result is a forward model containing three major components. A grid of dipoles, representing discrete sources of recorded activity, an electrode location matrix describing the position of each electrode on the head, and a lead field matrix, which transforms a signal generated at the dipole level to the signal recorded at the electrode level. This forward model was used to generate the ground truth signal as described below.

##### Generate a source signal

The initial ground truth signal was a linear sinusoidal chirp. Its frequency increased from 2Hz to 30Hz over 1 second, sampled with a resolution of 1000 Hz, mirroring the sampling rate of the EEG recording. The amplitude was constant and equivalent to 2 *µV* at the source level. The chirp was generated by adapting code from Cohen (Cohen, 2017). We selected a chirp for two related reasons. Firstly, the time and frequency dynamics of event-related potential (ERPs) vary from component to component. Thus, we wanted to use a signal that would encompass a large variety of possibilities. Secondly, this project has a methodological focus, so we did not want to limit the analyses to one specific component.

##### Projecting the signal to sensor space

After generating the forward model and the chirp, we utilized the Simulating Event-Related EEG Activity (SEREEGA) toolbox (Krol et al., 2018) to project the chirp signal to the electrode space. We chose a dipole near the parieto-occipital cortical surface (X = 25, Y = -80, Z = 25, MNI coordinate space) as the source of the ground truth signal. As EEG recordings depict the activity of pyramidal neurons oriented perpendicularly to the scalp (Nunez & Srinivasan, 2006), we constrained the dipole’s orientation to be perpendicular to the scalp surface. Modelling the electrode activity involved a linear combination of the lead field and the dipole signal. Specifically, SEREEGA solves the equation (refer to section 2.1 in Krol et al. (2018)):

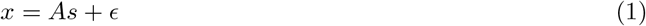

Here, *A* is the lead field matrix from the forward model, *s* is the chirp signal at the source level, and *E* is a noise vector excluded from our model. The lead field was normalized to ensure the signal’s amplitude at the electrode level reflected that at the source level, providing full control over the simulated signal. *x* is an *n* by *t* matrix representing the chirp signal of length *t* as recorded by the *n* EEG electrodes on our model’s scalp. The signal contained in this matrix is what we called ground truth, because it represents a known neuronal signal as it would have been recorded by our EEG system.

##### Combining EEG and ground truth

The ground truth signal lacks noise components and needs to be added to a real EEG recording. Following (Urigüen & Garcia-Zapirain, 2015) we term this process *injection*. To execute the injection, we simulated 300 events. Event onsets were selected at random to ensure that they did not contain any systematic activity associated with performing the Simon task. With the ground truth signal lasting 1000 ms and the longest baseline period also 1000 ms, we enforced a minimum 3000 ms delay between consecutive events. For any given event, the injection of the ground truth signal can be simply described as:

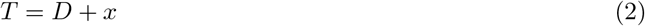

Here, *x* represents the ground truth matrix obtained earlier, *D* is the matrix representing the EEG recording time-locked to one event, and *T* is the resulting trial data.

This process yielded an EEG recording comprising 300 trials with a known ground truth signal time-locked to the trial onset. While a perfectly time-locked activity is not a realistic assumption, we refrained from jittering the chirp onset relative to the event onset to provide the pipelines with the best-case scenario. Because of the perfect time-locking, we can ensure that any distortion to the ground truth after preprocessing can be attributed to the preprocessing steps. Importantly, the maximum amplitude of the ground truth signal (2.208 *µV*) was extremely low compared to the amplitude of the raw EEG data (*Mdn* = 2000 *µV*).

#### 1.1.2 Pipeline selection

We tested six published or publicly available pipelines. Here we report the name and the associated sources, while a detailed description is reported in the supporting documentation online https://osf.io/rg5ah/files/osfstorage. Note that we named the pipelines according to their published name or by their main author. To avoid confusing authors with pipelines, throughout the paper we use italics to refer to the pipeline.

- *EEGLAB*: Automated ERP pipeline reported on the [EEGLAB website](https://eeglab.org/tutorials/11_Scripting/automated_pipeline.html#running-an-erp-pipeline).
- *Delorme_2023* : Pipeline optimised for event-related potentials (ERPs) described by Delorme (2023). Code is available [here](https://github.com/sccn/eeg_pipelines/blob/master/eeglab/process_eeglab_template.m).
- *Makoto*: Pipeline based on “Makoto’s preprocessing pipeline” [webpage](https://eeglab.ucsd.edu/wiki/Makoto%27s_preprocessing_pipeline).
- *Prep*: Automated pipeline described in Bigdely-Shamlo et al. (2015). This is an early-stage preprocessing pipeline, meaning that it performs a subset of cleaning procedures and researchers can expand it as they need. The reader should note that, although we refer to the pipeline as Prep, we have included extra steps, like ICA, which are not part of the original publication. Consequently, this is our version of *Prep*, based on existing practices in some laboratories of our department.
- *Henare_2018* : This is a pipeline previously used in our laboratory. We named from the first author of the preprint we used to implement it (Henare et al., 2018).
- *Henare_2018_Once*: this is a variation of *Henare_2018* where we removed the application of ICA on a copy of the EEG dataset filtered at 1 Hz.

We selected these pipelines based on practices followed in our offices, limiting the choice to automated pipelines implemented for EEGLAB. We aimed to implement each pipeline as closely as possible to the original, using provided scripts or following their associated information. In a few instances, we made minor changes to ensure the pipelines could run on our data without issues. Any deviation from the original scripts is highlighted in the supporting documentation online https://osf.io/rg5ah/files/osfstorage.

#### 1.1.3 Analyses

To evaluate the pipelines, we developed an adversarial approach where any two pipelines are compared by calculating the probability that one pipeline returns a signal closer to the ground truth than the other. In essence, given pipelines *P*_*A*_ and *P*_*B*_, we utilized a permutation-based approach to compute the probability of *P*_*A*_ returning a signal more similar to the ground truth than *P*_*B*_. Starting from the preprocessed data, the process involves the following steps:

1. Extract only the epochs common to all pipelines.
2. Trim the epochs by removing the baseline period. The final epochs only contain the injected signal.
3. For each pipeline, randomly sample and average N epochs to obtain grand average waveforms.
4. Compute the root mean squared error (RMSE) between each grand average and ground truth signal.
5. Repeat steps 3 and 4 *I* times, storing the RMSE values at each iteration.
6. For each combination of two pipelines (*P*_*A*_, *P*_*B*_), compute how many times, out of *I* repetitions, *P*_*A*_ RMSE values were lower or equal to *P*_*B*_ values.

We will discuss each step separately.

*Step 1*. During EEG preprocessing it is common to lose some data. Consequently, each pipeline will return only a subset of the original trials. To ensure that the comparisons are not biased by the specific trials used, we performed analyses only on the subset of trials that were retained by all pipelines.

*Step 2*. The ground truth chirp lacks a baseline period; therefore, we restrict the adversarial analyses to the period where the signal is expected.

*Step 3*. Averaging across trials is a simple way to eliminate non-systematic noise. The number of trials included in the grand average (*N*) varies from study-to-study, so here we set it to 100, an arbitrary yet realistic value. For each permutation, we used the same N trials across all pipelines to ensure that the grand averages were computed using the same portions of data.

*Step 4*. The RMSE for a pipeline *P* was computed as:

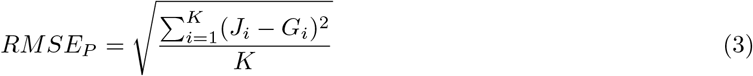

Where *J*_*i*_ represents the *i* sample of the grand average for pipeline *P, G*_*i*_ represents the equivalent sample of the ground truth signal, and *K* is the number of samples making up the signal. The value of *K* is determined by the sampling rate. The pipelines either preserved the original sampling rate (1000 Hz) or downsampled the data to 250 Hz. To ensure accurate RMSE computation, we created a downsampled copy of the ground truth signal at 250 Hz. Each pipeline was then compared against the ground truth sampled at the same rate. RMSE values closer to 0 indicate that the ground truth is close to the grand average, representing a pipeline that was capable to denoise the data without impacting the signal of interest. Higher RMSE values would indicate residual noise impacting the grand average, distortion of the injected ground truth during preprocessing, or a combination of both. Thus, the RMSE serves as a similarity measure simultaneously capturing both incomplete noise reduction and signal distortion effects.

We limited the computation of the RMSE to only one electrode. Specifically, we selected the electrode that showed the strongest amplitude in the ground truth data (PO4, max absolute amplitude = 2.208 *µV*). We assumed that this represents the best-case scenario for the pipelines. If a pipeline’s performance is suboptimal on the electrode with the strongest signal, improvement on weaker electrodes is unlikely.

*Step 5*. We repeated steps 3 and 4 100,000 times (*I* =100000). For each iteration, a new random sample of *N* trials was drawn to ensure that the RMSE is not biased by the specific trials included in each sample.

*Step 6*. At the end of the iterations, each pipeline had *I* RMSE values. To compare two pipelines, we calculated the probability that the RMSE values of one pipeline are lower than or equal to the RMSE values of another one:

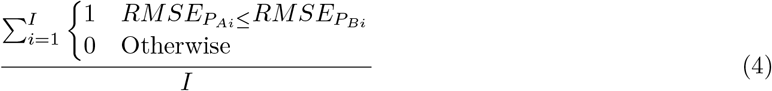

Where 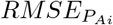 is the *i*th RMSE value for pipeline *A* and 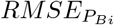 is the RMSE value for pipeline *B*, and *I* are the number of permutations performed.

##### Trial number effect

The previously described results are contingent on the number of trials used to compute the grand averages. We pre-set this number to 100 trials to represent an arbitrary but realistic value for ERP research. However, it is possible that the performance of each pipeline varies as a function of the number of trials averaged. To account for this possibility, we replicated the analyses described above, while systematically adjusting the number of trials used to obtain grand averages, ranging from *N* = 1 to *N* = 225 (the latter represents the maximum number of trials common to all pipelines).

#### 1.1.4 Results

We compared the efficacy of the pipelines through a *permutation-based adversarial approach*. Each pair of pipelines was tested by computing the probability that one pipeline returned a signal more or equally similar to the ground truth compared to another pipeline (see Analyses section). We would like to remind the reader that the comparisons are idiosyncratic to the data, ground truth and pipelines tested and can vary from study to study, reflecting the fact that different pipelines could be more adequate in different contexts. The results of this method are descriptive by design to refrain from interpreting the comparisons with a definitive black-and-white statement.

##### Fixed trial number

Figure 2 (I) reports the results of the adversarial analyses when 100 trials are used to compute grand averages. No pipeline consistently returns RMSE values lower than the others, evidenced by the fact that no pipeline showed values of 1 across all comparisons. We can explore specific trends by looking at the adversarial analyses within one pipeline. For instance, *Henare_2018_Once* and *Henare_2018* consistently exhibit probabilities exceeding 0.7 to produce RMSE values lower or equal to *Delorme_2023, Makoto*, and *Prep*. However, *EEGLAB* has a similar probability to produce a signal equivalent to or better than *Henare_2018_Once* and *Henare_2018*. Conversely, *Delorme_2023* and *Makoto*’s pipelines display a low probability of returning RMSE values lower or equal to the others. *Delorme_2023* yielded probabilities at or below 0.15 of returning RMSE values lower or equal to the others’ across all comparisons, while *Makoto* exceeded 0.5 only against *Delorme_2023*.

**Figure 1.**
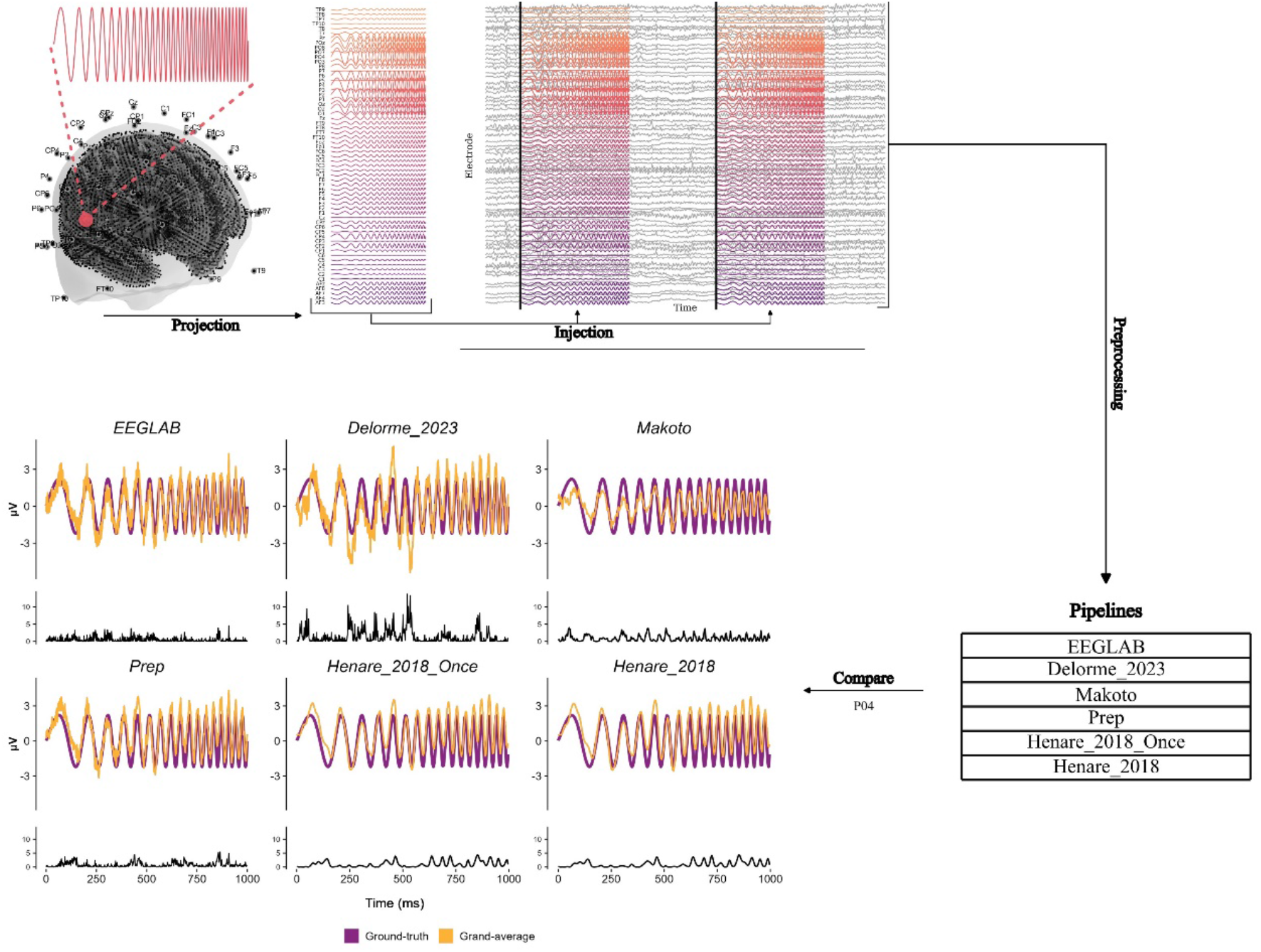
Projection. A forward model was utilized to project a chirp signal from a brain source (red circle) onto the scalp, resulting in a matrix representing the chirp as recorded by EEG electrodes. This matrix served as the ground truth activity for each electrode. **Injection**. The ground truth matrix was introduced into a real EEG recording at the onset of simulated events. These events were randomly placed across the EEG recording to create trials with known time-locked ground truth activity. Consecutive trials could not overlap. **Preprocessing**. The injected dataset underwent processing using six different pipelines, all of which were previously published or publicly available. **Compare**. For each pipeline, a sample of clean epochs was averaged (yellow waveforms). For this illustration, we randomly selected 100 trials to compute the grand averages. The effects of a pipeline were assessed by calculating the Residual Mean Squared Error (RMSE) values between each average and the ground truth signal (purple waveform) at electrode PO4. We selected PO4 because it contained the ground truth signal with the highest amplitude. The RMSE values were then entered into a permutation-based adversarial analysis process to obtain the probability of each pipeline providing lower or equal residuals to any other one. The black lines underneath each plot represent the squared residuals between each grand average and the ground truth at each time point.

**Figure 2.**
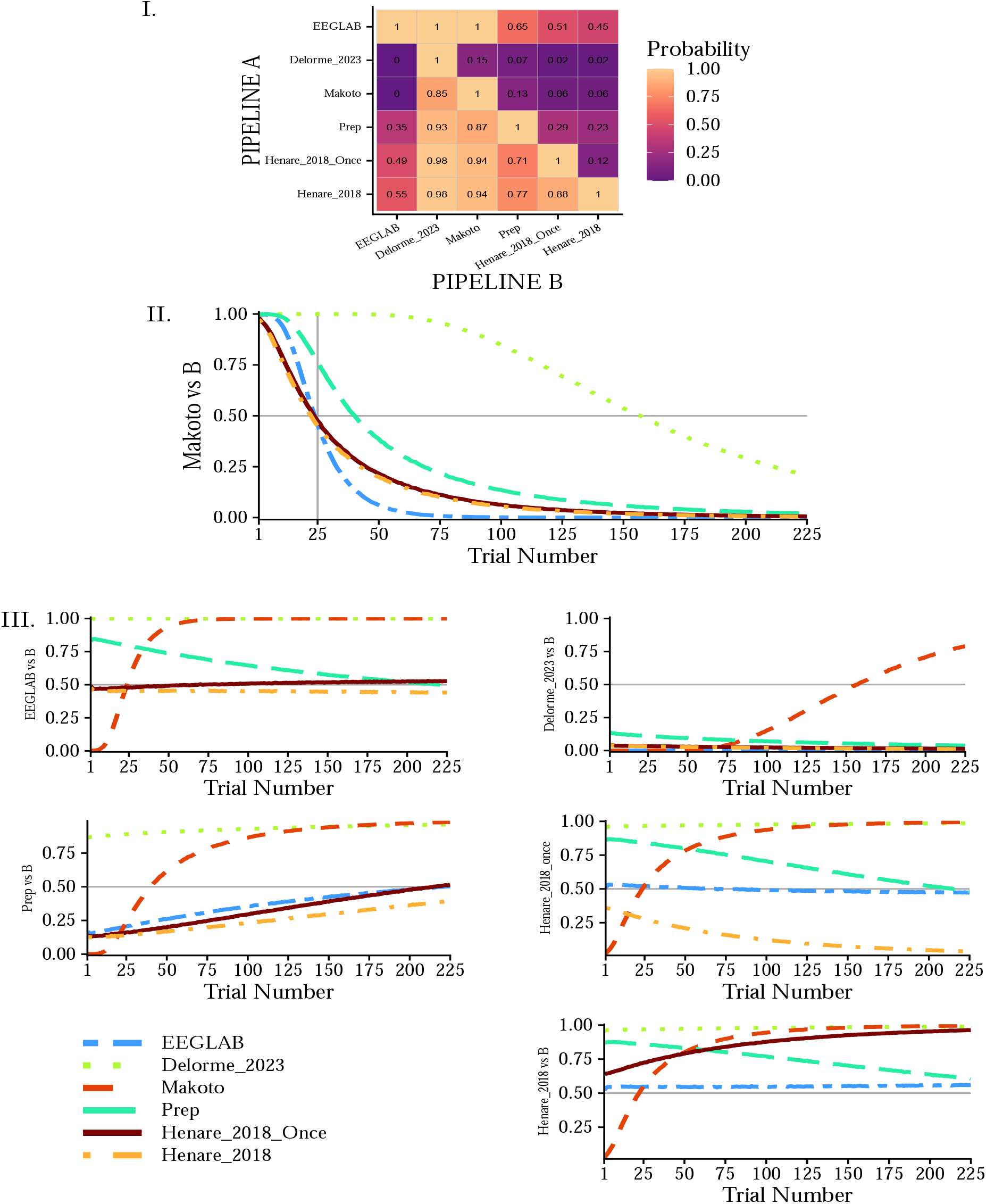
Adversarial analysis results. **I.**Probability matrix showing how often PIPELINE A yields lower or equal RMSE than PIPELINE B, based on grand averages relative to the ground truth. Each cell can be read as “PIPELINE A has probability p of being less or equally distorted than PIPELINE B” (e.g., *Prep* vs. *EEGLAB*: 0.35). Values are rounded to two decimals. **II**. Adversarial comparison between *Makoto*’s pipeline and all others as a function of the number of averaged trials. Each line represents the probability that *Makoto*’s pipeline achieves lower or equal RMSE. This probability decreases as more trials are averaged; with fewer than 25 trials, *Makoto*’s pipeline outperforms all others (all probabilities > 0.5). **III**. Same analysis as in *II*, but shown for all other pipelines. Overall, pipeline comparisons depend strongly on whether and how many trials are averaged in the analysis.

**Figure 3.**
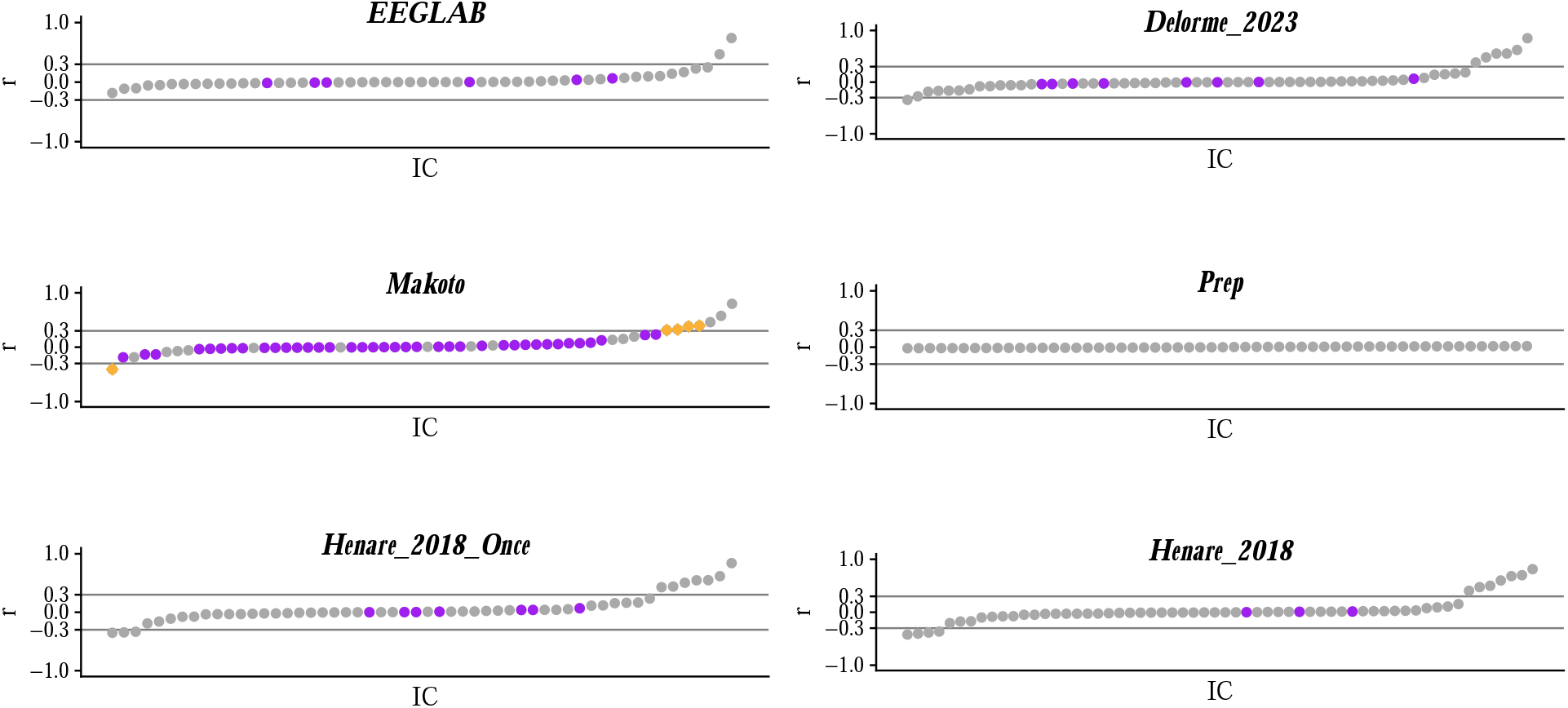
Correlations between the chirp ground truth and the ICs ordered from low to high values. Gray dots represent ICs preserved. Purple dots represent ICs removed from the data. Yellow dots represent ICs that were removed from the data but were highly correlated (r ≤ 0.3 or r ≥ 0.3) with the ground truth. Note that in Study 1 we did not correct for this, meaning that pipelines were able to remove highly correlated ICs. Only *Makoto*’s pipeline showed this behaviour, removing 5 ICs that likely contained the injected chirp signal. *Makoto*’s pipeline retains only ICs that are considered representing brain activity, meaning that in this case the chirp was not only isolated through ICA (high correlations), but also flagged as a non-brain signal. This realisation prompted the second study.

##### Trial number effects

While the results above suggested that *Delorme_2023* and *Makoto*’s pipelines are less likely than other pipelines to return a chirp closer to the ground truth, Figure 2 (II) provides a more nuanced perspective. *Makoto*’s pipeline displays a clear decreasing trend, where the probability to provide lower or equal RMSE values decreases with the increase of the number of trials used to compute grand averages. When the number of trials is limited (approximately ≤ 25), *Makoto*’s pipeline consistently exhibits probabilities exceeding 50% to produce a signal closer to the ground truth (it performs better) across all comparisons, except *Prep* and *Delorme_2023*, where this threshold is reached with higher number of trials.

Figure 2 (III) shows the trial effect for all the remaining pipelines. We observe different trends for each comparison. For instance, while *Prep* probabilities tend to increase with the trial number, the probabilities of *Delorme_2023* remain stable (with the exception of *Delorme_2023* vs *Makoto* as discussed above). Similarly, the adversarial comparisons for *EEGLAB* vs *Henare_2018_Once* (*P* (*RMSE* _*EEGLAB*_ ≤ *RMSE*_*Henare*_2018_*Once*_): *M* = 0.507, *SD* = 0.017, Range: 0.468;0.529) and *EEGLAB* vs *Henare_2018* (*P* (*RMSE* _*EEGLAB*_ ≤ *RMSE* _*Henare*_2018_): *M* = 0.450, *SD* = 0.005, Range: 0.440;0.479) produces similar probabilities independently of the trial number. The interested reader can systematically explore the results in a dedicated app.

##### Descriptives

Pipelines vary in the time and resources they require, an aspect researchers might consider when selecting how to clean their data. We recorded the time each pipeline took to preprocess our dataset (Table 1). All pipelines took less than 10 minutes, except for *Makoto*’s pipeline, which took over 15 minutes (1026 seconds), and *Prep*, which ran for over an hour (5232 seconds). We also recorded the number of channels removed and interpolated, as well as the number of independent components (ICs) removed from the data, as shown in Table 1. All pipelines removed and interpolated less than 15% of the electrodes (*MIN* = 2, *MAX* = 9). Regarding the ICs, *Makoto*’s stringent criteria led to the removal of 43 ICs. Although all the other pipelines used the same set of exclusion criteria, they removed varying numbers of ICs, with *Prep* excluding none and *Delorme_2023* removing 8.

**Table 1:** Pipelines run time, interpolated channels and removed ICs.

| Pipeline | Computation Time [s] | Interpolated Channels | IC Removed |
| --- | --- | --- | --- |
| *EEGLAB* | 333 | 9 | 6 |
| *Delorme_2023* | 406 | 2 | 8 |
| *Makoto* | 1026 | 4 | 43 |
| *Prep* | 5253 | 6 | 0 |
| *Henare_2018* | 348 | 2 | 3 |
| *Henare_2018_Once* | 301 | 8 | 7 |

*Note.* Computation time, number of interpolated channels, and number of removed ICs for each pipeline can be considered alongside the results of the adversarial analyses when selecting a pipeline. For instance, one might need a fast preprocessing pipeline or one that removes the least amount of data.

### 1.2 Interim discussion, limitations and second iteration

The probability matrix and trial effect results highlight the nuanced view provided by our method. Each pipeline exhibits specific trends in the adversarial analyses that can only be understood within the broader context of all the comparisons investigated. Just because one method scores a high probability of producing lower or similar RMSE than another does not mean it does so consistently across all comparisons. Moreover, we showed that a pipeline’s ability to return a signal close to the ground truth can depend on the number of trials averaged, if any, after data cleaning. Consequently, researchers developing a preprocessing script should consider the number of trials available and whether they will be averaged. In our case, if 100 trials per condition are available, we might opt for *Henare_2018* or *EEGLAB*, especially if we want to use a publicly available pipeline. However, if we have only access to a small number of trials per condition, *Makoto*’s pipeline may be preferable.

A significant caveat regarding the adversarial results obtained here is that they may be biased by the way each pipeline applied ICA to remove noise from the data. We initially did not consider that ICA might isolate the ground truth signal, leading to its associated component being removed from the data. Essentially, the ground truth could be mistaken for noise and eliminated. This scenario could occur in all pipelines, except *Makoto*’s, exclusively through `IClabel` (Pion-Tonachini et al., 2019). `IClabel` estimates the likelihood of each IC representing brain activity, specific types of noise (eye, muscle, cardiac activity, line noise), or something not specified. Any IC having a probability of representing noise higher than a user-defined threshold is then removed from the data. In our case, *EEGLAB, Delorme_2023, Prep, Henare_2018_Once* and *Henare_2018* only discarded ICs with a 90% probability or higher of reflecting eye or muscle noise. *Makoto*’s pipeline adopted a more stringent approach, retaining ICs only if:

1. They exhibited a higher probability of representing brain activity compared to any other type of activity, as determined by `Iclabel`.
2. Their associated dipoles were located within the brain.
3. The IC scalp maps of their associated dipoles had a residual variance lower than 15%.

In the first study, if an IC associated with the ground truth failed any of these tests, it would have been removed from the data.

Considering this, we hypothesised that *Makoto*’s pipeline was more likely to return equal or lower RMSE at a low number of trials because of its tendency to aggressively remove data from the recordings. By removing considerably more activity after ICA than other pipelines, it might have removed more noise but also more signal associated with the chirp. When not many trials are averaged, this might be advantageous, as computing grand averages is not possible (single-trial case) or not as effective. However, this approach might be counterproductive when more trials are averaged. In that case, a better strategy could be to retain more noise and more signal, assuming that the residual noise will average out at the end of preprocessing.

To ensure that our comparison between pipelines with different ICs exclusion/inclusion criteria was not unfairly favouring some pipelines over others, we conducted a second study that employed the same analyses, a variation of the same pipelines, and a new ground truth.

### 1.3 Study 2

Here we aimed to verify that the initial results were not predominantly influenced by a pipeline decomposing the ground truth and labelling it as noise during the ICA + `IClabel` step. To solve this problem, we aimed to remove any unfair advantage/disadvantage between pipelines during ICs removal while maintaining the original variability in criteria between them. To achieve this, we took two steps:

1. Generate a new ground truth that, if decomposed during ICA, had higher probability to be labelled by `IClabel` as representing a brain signal
2. Enforce all pipelines to retain those ICs that were likely to contain the ground truth, even if the pipeline’s default parameters would have removed them

In the following sections we will explain these two points in more details.

#### 1.3.1 Dataset Generation

##### Generate ground truth signal

To minimise the likelihood of the ground truth being isolated and removed by ICA, we needed a signal that was highly likely to represent brain activity. To identify such a signal, we collected a new dataset from the same author (DS) performing a 2-dimensional mental rotation task of letters (Cooper, 1975). The data was filtered using a non-causal one-pass zero-phase finite impulse response filter with a Hamming window implemented in `pop_eegfiltnew` by EEGLAB. Specifically, we applied a high pass filter (cutoff frequency = 1 Hz at -6 dB, transition bandwidth = 2 Hz, passband edge = 2 Hz, filter order = 1651 points) and a lowpass filter (cutoff frequency = 22.5 Hz at -6 dB, transition bandwidth = 5 Hz, passband edge = 20 Hz, filter order = 661 points). We chose these parameters to avoid unintentionally biasing the results, as they were not employed by any pipeline. The sections of data before and after the experiment were removed to reduce noise, and ICA was applied to this lightly processed dataset. For this step, we used the Jader algorithm (Cardoso & Souloumiac, 1993), again, because it was not used by any of the pipelines in our study. Finally, we segmented the data into 300 ms epochs starting from the stimulus onset. Given that the sole purpose of this dataset was to provide a more realistic ground truth signal, no trials were rejected, and we did not differentiate between experimental conditions.

We applied `IClabel` to categorise the extracted ICs, and their source dipoles were estimated through `dipfit`, using default parameters. Among all ICs, we selected one that met the following criteria:

1. It was considered to represent brain activity according to `IClabel`
2. Its associated dipole was inside the brain
3. The scalp map of its associated dipole had a residual variance lower than 15%
4. It did not appear to capture only noise according to visual inspection

Notably, the first three criteria were the same as those employed by *Makoto*’s pipeline. We made this choice to ensure that we identified a ground truth signal that would be considered brain activity even by the most stringent pipeline. Visual inspection was included because the dataset was only filtered before ICA, making the resulting ICs likely to be noisy.

Once we identified an appropriate IC, we extracted its scalp activity and averaged it across all epochs. The final result was an *n* by *t* matrix representing the scalp activity of the IC at *n* electrodes over *t* samples. This matrix represented the new ground truth.

##### Combining EEG and ground truth

We injected the ground truth signal following the same procedure and dataset as Study 1 (see Study 1 Projection section). The ground truth was injected at the onset of the same 300 events used in the first study, so that the only difference between studies was the ground truth signal itself.

##### Preprocessing

We used the same pipelines as in study 1 (Pipeline selection). However, we took an extra step to ensure that the ground truth signal would not be entirely removed through ICA. After ICs were extracted, we created a duplicate of the data. This duplicated data was then segmented around the simulated events, spanning from the event onset to 300 ms afterwards, equivalent to the duration of the ground truth signal. This segmentation allowed us to derive the activity of each IC when the ground truth was expected. For each IC, we averaged its scalp activity across all the epochs, and we calculated the linear correlation between the averaged IC and the ground truth activity. Any component exhibiting a moderate to high correlation with the ground truth (*r* <0.3 or *r* >0.3) was retained regardless of whether it met the inclusion criteria set by each pipeline. This step was added to every pipeline.

#### 1.3.2 Analyses

We conducted the same analyses described in Study 1, computing the probabilities of each pipeline to provide a better or equally good result as any other and by assessing the trial number effect.

#### 1.3.3 Results

We first verified the number of ICs that were forced to be retained through the correlation procedure described above. Figure 4 shows that the correction was unnecessary for all pipelines except *EEGLAB* and *Delorme_2023*. These two pipelines flagged one IC each (in yellow) for removal, even if the IC was likely to contain the ground truth signal (*r* > 0.3). Because of the correction applied in this study, these components were retained in the final data.

**Figure 4.**
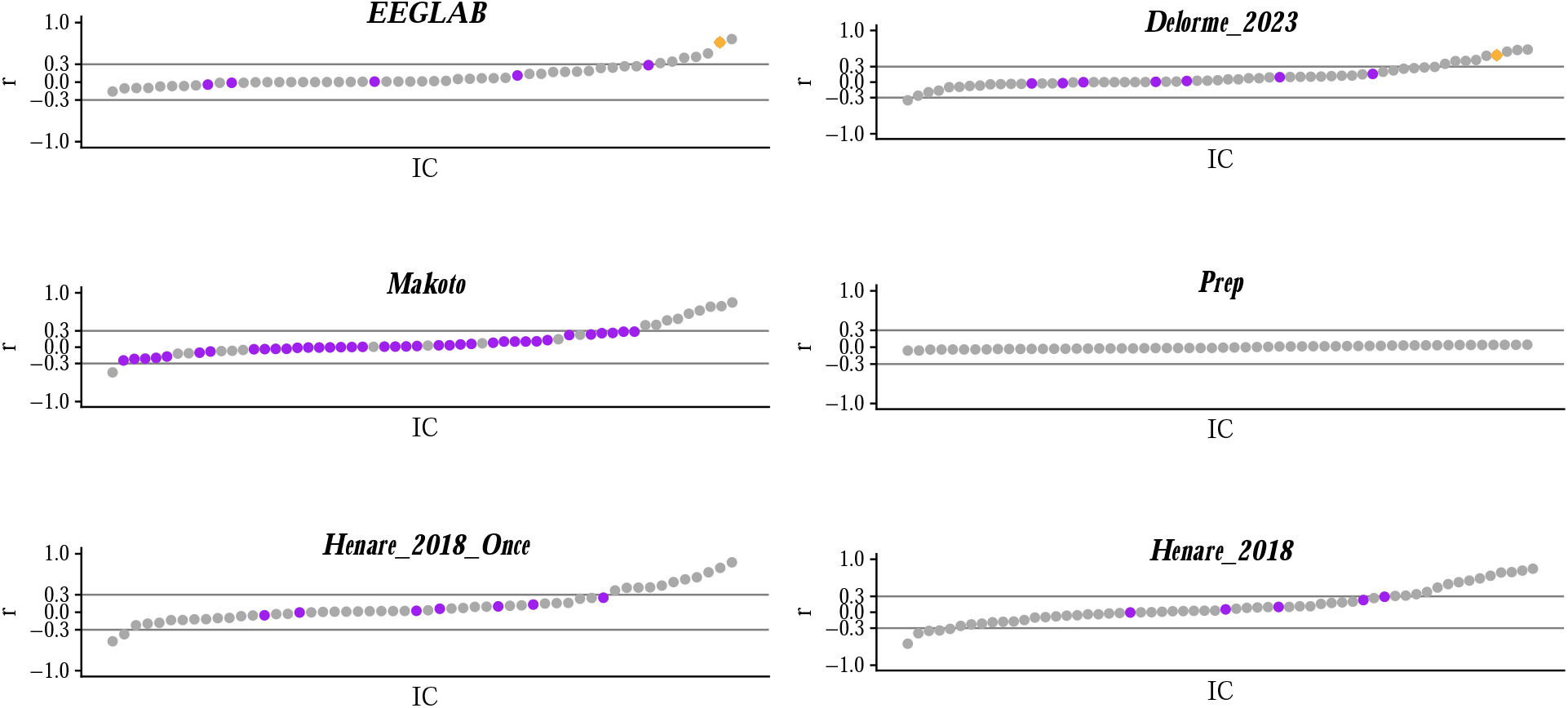
Correlations between ICs and the ground truth ordered from negative to positive values. Gray dots represent ICs preserved in the data, purple dots represent ICs removed by the pipeline, and yellow dots represent ICs flagged for removal but preserved due to high correlation with the ground truth (*r* ≤ -0.3 or *r* ≥ 0.3). Most pipelines show ICs highly correlated with the ground truth, but only *EEGLAB* and *Delorme_2023* considered one of these ICs each as noise. These results suggest that the new injected signal, even if decomposed during ICA, was not deemed relevant noise by `IClabel`. Notably, *Makoto*’s pipeline only flagged and removed low-correlated ICs indicating that it considered all highly correlated ICs as representing intrinsic brain activity.

##### Fixed trial number

Overall, the probability matrix yields similar results to Study 1 (Figure 5) (I). *Henare_2018* and *Henare_2018_Once* show probabilities higher than 0.7 of achieving higher or equally good RMSE values across all comparisons, even against *EEGLAB*, which in study 1 showed a similar performance. *Prep* demonstrates an increase in the probability of achieving higher or equally good RMSE values compared to *EEGLAB*, rising from 0.353 to 0.629. *Delorme_2023* and *Makoto*’s pipelines are still less likely to provide lower or similar RMSE values across all adversarial analyses. However, *Delorme_2023* now has a higher probability than *Makoto*’s pipeline to provide a signal closer to the ground truth (*P* (*RMSE*_*Delorme*_2023_ ≤ *RMSE*_*Makoto*_) = 0.68).

**Figure 5.**
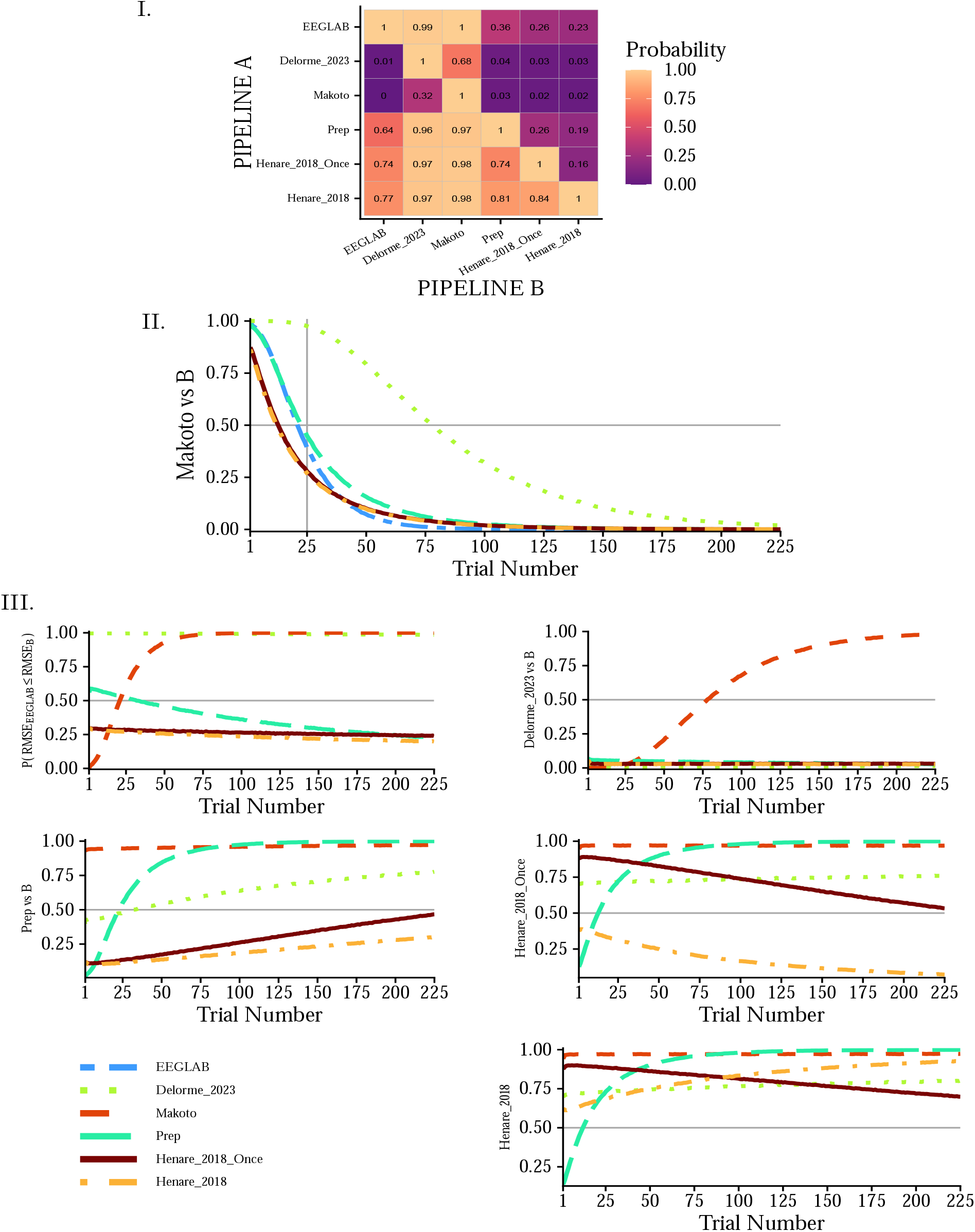
Adversarial analyses results for the new ground truth signal. Plots are equivalent to those reported in Figure 2. **I.**Although we used a different ground truth and we ensured that no pipeline removed most of it during ICA, the probability matrix shows similar patterns as in Study 1. **II**. *Makoto*’s pipeline still shows a clear trial-number effect, with high likelihood of returning lower or equal RMSE values than any other method when a low number of trials is averaged. **III**. Similarly to *II*, every other pipeline shows trends like those obtained in the first study.

##### Trial number effects

As shown in Figure 5) (II), the inverse relationship between probability and trial number persists for *Makoto*’s pipeline. The likelihood of obtaining results closer or equal to the ground truth when using *Makoto*’s pipeline is higher than 0.5 across all adversarial comparisons when the number of trials used to create the grand averages is low. Echoing the observations obtained using 100 trials, from Figure 5) (III) we can see that *EEGLAB* has a stable probability below 0.5 against *Henare_2018_Once* (*P* (*RMSE*_*EEGLAB*_ ≤ *RMSE*_*Henare* _2018_ *Once*_): *M* = 0.265, *SD* = 0.013, Range: 0.244; 0.299), and *Henare_2018* (*P* (*RMSE*_*EEGLAB*_ ≤ *RMSE*_*Henare* 2018_): *M* = 0.232, *SD* = 0.024, Range: 0.198; 0.294), while its likelihood of providing higher RMSE values against *Prep* decreases with the increasing number of trials (*P* (*RMSE*_*EEGLAB*_ ≤ *RMSE*_*Prep*_): *M* = 0.374, *SD* = 0.102, Range: 0.241; 0.602).

##### Descriptives

Run times, number of interpolated channels, and number of ICs removed are reported in Table 2. The time each pipeline took to preprocess the dataset was similar to the times observed in Study 1 (Table 1), with the exception of *Prep*, which ran for about half a hour (1709 s). Similarly, the number of channels and ICs removed was on par with that observed in the first study, possibly indicating reliability in the pipelines’ ability to detect noisy channels and noise-associated ICs.

**Table 2:** Pipelines run time, interpolated channels and removed ICs.

| Pipeline | Computation Time [s] | Interpolated Channels | IC Removed |
| --- | --- | --- | --- |
| *EEGLAB* | 565 | 9 | 5 |
| *Delorme_2023* | 356 | 2 | 7 |
| *Makoto* | 1306 | 4 | 38 |
| *Prep* | 1802 | 6 | 0 |
| *Henare_2018* | 359 | 1 | 5 |
| *Henare_2018_Once* | 315 | 8 | 7 |

*Note.* Computation times, number of interpolated channels and number of removed ICs are similar to those observed in the first study. The major difference is in the time the Prep pipeline took to run, which went from 5232 to 1802.

### 1.4 Interim discussion

In the second study, we verified that our results were not solely driven by the criteria each pipeline employed to retain ICs after applying ICA to the data. This step was necessary because one of the pipelines included in our test (*Makoto*) had stricter ICs inclusion criteria compared to all the other ones. By observing the correlation values between ICs and the ground truth signal, we can see that every pipeline, except for *Prep*, likely decomposed the ground truth into one or more components. Only the *EEGLAB* and *Delorme_2023* pipelines flagged one of these components as being noise, indicating that the new ground truth signal was usually considered as representing intrinsic brain activity. Notably, this was true even under the most stringent criteria employed by *Makoto*’s pipeline.

Because the ground truth was itself an IC that should have reflected brain activity, it is reassuring to see most of the pipelines consider it in the same way, even if the signal was extracted from a different dataset. However, the behaviour of *EEGLAB* and *Delorme_2023* pipelines is less obvious. Because of the `IClabel` settings they implement, these two pipelines must have recognised one component highly correlated with the ground truth as representing muscle or eye noise. We can look at these results in two ways. On the one hand, if the ground truth really captured only brain activity, then the flagging of these components is a false alarm. Because *Henare_2018* and *Henare_2018_Once* used the same `IClabel` settings but did not recognise any highly correlated component as representing muscle or eye noise, it is possible that the cleaning steps preceding ICA and *IClabel* influenced the labelling behaviour. On the other hand, if the ground truth we used did not really capture brain activity, then every other pipeline would wrongly consider an exogenous noise signal as endogenous brain activity, a scenario that would raise questions regarding ICA and automatic labelling of ICs. Although both scenarios are possible in theory, we believe the first one to be more plausible because of the way the ground truth was obtained and, as we will discuss now, because the results are similar to those obtained in Study 1.

When 100 trials were averaged together, the adversarial analyses showed the same trends we observed with the simulated chirp, suggesting that the comparisons are not only driven by the way each pipeline uses ICA. This conclusion is also supported by the qualitative similarity across studies in the results obtained by systematically modifying the number of trials averaged before computing the RMSE values. Moreover, even the descriptive measures showed only minor changes, indicating that the behaviour of the tested pipelines was generally unaffected by the ground truth we used.

In conclusion, the second study validated the reliability of the methodology and results obtained in Study 1. We showed that the original results were not fully driven by the pipelines being able to recognise the injected signal as non-brain activity. Importantly, the step introduced to avoid the removal of possible ICs associated with the ground truth (used only for two *EEGLAB* and *Delorme_2023*) was necessary only because of the variability in inclusion and exclusion criteria between pipelines. If one were to test pipelines that employ the same IC retention rules, this step might be unnecessary. In that case, similar ICs should be removed by the different preprocessing methods, without a drastic difference like the one observed between *Makoto*’s pipeline and the others tested here. Moreover, if a difference was to be observed, then it could be investigated to understand which preprocessing steps influence the removal and retention of ICs.

### 1.5 Discussion

A significant amount of effort has been devoted to identifying the best methods for cleaning and preparing EEG data (Bailey, Hill, et al., 2023; Bailey, Biabani, et al., 2023; Fló et al., 2022; Gabard-Durnam et al., 2018). However, researchers often need to modify prepackaged pipelines, tweak parameters, or create custom pipelines tailored to their specific studies. In this paper, we introduced a flexible framework to assist researchers in making informed decisions about preprocessing pipelines and parameters. Our method employs a realistic head model to generate a ground truth signal at the cortical level, which is then injected into real EEG data containing natural experimental noise. By preprocessing this dataset with various pipelines, we assessed their ability to remove noise without distorting the signal of interest using the RMSE between the processed signal and the ground truth. Additionally, we implemented a resampling-based adversarial approach to compare each pair of pipelines, resulting in a probability matrix that indicates the likelihood of one pipeline performing equally well or better than another.

In a series of two studies, we observed that, perhaps unsurprisingly, the performance of any pipeline depends on the number of trials used to obtain grand average waveforms after preprocessing. When the number of averaged trials was low, the best performing pipeline across all adversarial comparisons was *Makoto*’s. However, the adversarial performance of this pipeline monotonically decreased as the number of trials averaged increased. Conversely, pipelines such as *Prep* and *Henare_2018*, showed increasing likelihood of returning a signal closer to the ground truth as the number of trials increased. These trends can be explained by considering that data cleaning procedures not only reduce or remove noise, but could also affect the signal of interest. Potentially, the more aggressive a procedure is, the more likely it will influence the signal of interest. When only a few trials are available, the benefits of removing as much noise as possible could be beneficial, even if the signal of interest is partially altered. However, with a higher number of trials, one can rely on averaging to remove non-systematic noise. That is, one could decide to retain more noise at the single-trial level. Averaging can then be exploited to return a clean and undistorted signal.

The results presented here are specific to the combination of dataset, ground truth signal, selected pipelines, and the settings applied. While this might be viewed as a limitation, we consider it a strength of our approach. Each study has idiosyncrasies and requirements that may lead researchers to tailor their preprocessing pipelines accordingly. Consequently, it is difficult to determine whether results from published analyses comparing methods or pipelines remain valid when applied to different data or adjusted parameters. Additionally, every comparison is inherently limited to the methods tested. Just because one pipeline performs best in one study does not mean it will do so in another one. Our framework addresses these concerns in two ways. First, it allows researchers to choose which pipelines and parameters to test, influenced by factors such as planned analyses, knowledge of the signal of interest, EEG system used, available time, resources, and coding skills. This ensures that comparisons are relevant to specific research settings. Second, our adversarial analyses avoid making absolute statements about the best overall pipeline. The probabilistic and descriptive nature of the results should be interpreted within the context of the study. For example, in our work, if conducting a standard ERP experiment with at least 100 trials per condition, we might select the *Henare_2018* or *EEGLAB* pipelines. For single-trial analyses, we would opt for *Makoto*’s. Interestingly, if we were not ICA enthusiasts, we would opt for *Prep*. This pipeline performed similarly or better than others, especially with a high number of trials, even if ICA was not utilised. This means that other steps implemented were sufficient to remove noise without altering the signal of interest.

An important question we aimed to address with this framework is how to make informed decisions about the best data cleaning methods without examining the data itself. Recently, (Delorme, 2023) evaluated the efficacy of various preprocessing methods by analysing the percentage of channels showing significant differences between two experimental conditions. While Delorme’s analytical framework is conceptually similar to ours, the two methods could benefit from each other. Our study focused on a single channel, whereas Delorme utilised information from all electrodes. By expanding the RMSE analysis to include all channels, we could obtain a more comprehensive measure of a pipeline’s cleaning ability. However, unlike Delorme, our method is blind to any experimental conditions. We believe that the selection of cleaning techniques should be independent of the results of interest to avoid choosing a preprocessing method solely for the purpose of achieving significant results. Therefore, applying a ground truth injection-based framework and considering results across all electrodes may be the most effective way to make informed decisions about pipeline selection.

To conclude, we have presented a framework for selecting preprocessing methods that is flexible and produces interpretable but not absolute results. The flexibility stems from the ability to model a ground truth signal at your discretion, considering the specific research question, available data, and background knowledge. Moreover, results can be extracted from the same dataset that will be analysed in the main research without the risk of “hunting for a significant result”, because this method is based on a simulated ground truth and fictitious trials. The interpretability and flexibility of the results arise from the probabilistic nature of the adversarial analyses presented here. Researchers are provided with an accessible metric, but they must make an informed decision from it. The probability matrices and the probability-by-trial-number plots offer an overview of every possible comparison and are meant to provide guidance, not replace the researcher’s best judgment. As stated multiple times, each study has its own requirements, and we need to consider this. Some researchers might have access to hundreds of trials, some might not, and others are interested in conducting both grouped and single-trial analyses. The number of electrodes and their coverage differ from one EEG system to another, and the population studied can also be relevant. Because the research space in the field of EEG preprocessing is almost infinite and continuously developing, it is probably better to rely on a flexible framework that offers informed and tailored guidance rather than a firm statement that can hardly be generalised.

### 1.6 Guidance, modifications and future directions

The method we introduced in this work is highly flexible and can be adapted or expanded to suit the needs of different projects. In this section, we highlight some crucial aspects to consider when using this method and briefly provide possible ways to expand it.

The most crucial element to address is whether a pipeline could be tailored to each participant’s dataset. The answer is a strict *NO*. Preprocessing should be considered a variable that needs to be controlled in a study. Since researchers are usually not primarily interested in assessing the effects of preprocessing on their data, this variable should be kept constant across participants and conditions. Failing to do so would add an extra dependent variable that would need to be accounted for during analyses, making it more difficult to interpret the data.

The next question is how to select the dataset for the injection. The first option we suggest is to use a pilot dataset. Pilot data is often collected at the beginning of a study. A pilot dataset is ideal for the selection of a pipeline for the whole study because: (1) it is usually recorded with the same EEG system, electrode caps, in the same location, and with the same protocols as the data to be analysed; and (2) it is not included in the main analyses. However, if pilot data is not available, one could take a random dataset from those collected for the study.

Related to this issue is the number of datasets to which this method should be applied. While we utilised a single dataset in this instance, an alternative approach could involve using multiple pilot datasets. Another option, inspired by the method suggested by (Ostlund et al., 2022) for selecting the parameters of the Fitting Of One Over F algorithm (Donoghue et al., 2020), would be to take a random sample from the data that is to be analysed. By doing so, one could verify that the adversarial analyses are consistent across datasets. If the results suggest different pipelines for different datasets, it might indicate systematic differences across the sample sets (e.g., level or type of noise, duration of recordings, etc…) that should be considered when developing a preprocessing procedure. Importantly, if a sample of recordings is used, we suggest researchers: (1) keep the same sample throughout the selection process, especially if multiple iterations of this method are used; and (2) use a random number generator to select the sample, and set a seed to ensure that the analyses can be replicated.

In terms of modifying and tailoring the framework, the landscape of possibilities is vast. One obvious option is to define a specific ground truth signal, incorporating prior knowledge where possible. For example, researchers studying Visual Evoked Potentials (VEP) could generate a source signal resembling a canonical VEP and project it to the scalp to obtain a study-specific ground truth. A more flexible option involves using a chirp similar to that we employed, but analysing how residuals change across time over the chirp signal. By doing so, one could gather insights into whether a pipeline affects lower and higher frequency components differently. Studies focusing on the P300 component, for instance, would prioritise pipelines that produce lower RMSE values at lower frequencies of the chirp.

The dipole’s location and orientation could also be chosen to reflect the spatial information of the neural generators of the signal of interest. Similarly, when structural MRI (MRI) and EEG are recorded from the same participants, the structural MRI could be used to create a subject-specific forward model, potentially improving the ground truth definition. In the absence of structural MRI, we strongly recommend using a standard head model for realistic ground truth signal creation.

Another approach is to modify the RMSE analyses used here to reflect those used in the study. For instance, one could compare the time-frequency decomposition of the preprocessed signal to that of the ground truth. The specific metric used to assess signal similarity may vary depending on the analysis, and we leave this choice to the expertise of each researcher. However, we suggest that any metric defined should be applicable to the permutation-based adversarial analyses defined here. Similarly, the parameters measured can be expanded to assess the effects of preprocessing on various features of the signal of interest. Instead of comparing RMSE values for the entire signal, one could investigate how preprocessed data differ in terms of specific attributes such as latency or amplitude.

Finally, this technique can be used to assess the effects of a single pipeline and potentially optimise its parameters to try obtaining a cleaner neuronal signal. For those uninterested in comparing different pipelines, it could still be worth evaluating whether the way they preprocess their data influences the final signal significantly. In such cases, researchers could just extract a distribution of RMSE values (steps one to five in the Analyses section), skipping the final comparison between different methods. The distribution of RMSE would provide information regarding the consistency of the cleaning process and its ability to clean the data without introducing distortions. If parameters need to be adjusted to obtain a higher quality signal, then the adversarial analysis can be carried out across different versions of the same pipeline.

